# DNA binding is rate-limiting for natural transformation

**DOI:** 10.1101/2024.06.06.597730

**Authors:** Taylor J. Ellison, Courtney K. Ellison

**Affiliations:** Department of Microbiology, University of Georgia, Athens, GA

## Abstract

Bacteria take up environmental DNA using dynamic appendages called type IV pili (T4P) to elicit horizontal gene transfer in a process called natural transformation. Natural transformation is widespread amongst bacteria yet determining how different factors universally contribute to or limit this process across species has remained challenging. Here we show that *Acinetobacter baylyi*, the most naturally transformable species, is highly transformable due to its ability to robustly bind nonspecific DNA via a dedicated orphan minor pilin, FimT. We show that, compared to its homologues, *A. baylyi* FimT contains multiple positively charged residues that additively promote DNA binding efficiency. Expression of *A. baylyi* FimT in a closely related *Acinetobacter* pathogen is sufficient to substantially improve its capacity for natural transformation, demonstrating that T4P-DNA binding is a rate-limiting step in this process. These results demonstrate the importance of T4P-DNA binding efficiency in driving natural transformation, establishing a key factor limiting horizontal gene transfer.

**Importance:** Natural transformation is a multi-step, broadly conserved mechanism for horizontal gene transfer in which bacteria take up exogenous DNA from the environment and integrate it into their genome by homologous recombination. A complete picture of the factors that limit this behavior remain unclear due to variability between bacterial systems. In this manuscript, we provide clear and direct evidence that DNA binding by type IV pili prior to DNA uptake is a rate-limiting step of natural transformation. We show that increasing DNA binding in antibiotic resistant Acinetobacter pathogens can boost their transformation rates by 100-fold. In addition to expanding our understanding of the factors that limit transformation in the environment, these results will also contribute to a deeper understanding of the spread of antibiotic resistance genes in relevant human pathogens.

## Introduction

Natural transformation has been demonstrated in upwards of 80 species (1), aiding in the spread of antibiotic resistance genes and mobile genetic elements across groups of unrelated microorganisms (2). There are three major steps required for natural transformation: 1) DNA uptake mediated by T4P, 2) DNA translocation into the cytoplasm, and 3) incorporation of exogenous DNA into the chromosome. T4P are broadly distributed nanomachines (3) that are composed primarily of major pilin subunits that are polymerized or depolymerized via the activity of ATP-hydrolyzing motors to extend or retract (4, 5). The exact role of T4P in DNA uptake was unclear until recently when it was directly shown that pili from diverse species including T4P from *Vibrio cholerae* (6, 7) and the gram-positive competence pili found in *Streptococcus pneumoniae* (8) and *Bacillus subtilis*(9) bind DNA to mediate DNA uptake.

*Acinetobacter baylyi* is the most naturally transformable (also referred to as naturally competent) species reported in the literature, reaching transformation frequencies of up to 50% when grown in standard lab media like lysogeny broth (LB)(10, 11). *A. baylyi* produces T4P constitutively (11), and this is one likely explanation for its high rates of natural transformation. However, species like *Acinetobacter* pathogens that can be induced for increased T4P synthesis upon surface-associated growth only reach natural transformation frequencies of ∼1 per 10,000 cells (12, 13). This study aims to identify factors contributing to the high transformation frequencies of *A. baylyi*.

T4P-DNA binding is essential to the first step of DNA uptake, and we hypothesized that it may be a major limiting step during natural transformation. Previous studies showed that minor pilins mediate DNA binding in other competent species including *Vibrio cholerae* (6) and *Neisseria spp*. (14), and we suspected that a minor pilin may likewise be involved in DNA binding in *A. baylyi*. Many minor pilins are essential to T4P synthesis, with the primary model in the field suggesting they form a complex that first primes the T4P assembly machinery to aid in the recruitment of the major pilin subunit to T4P machines (15, 16). This prediction suggests minor pilins assemble in a T4P tip-associated complex that is essential for T4P synthesis. *A. baylyi* encodes six minor pilins in a conserved minor pilin operon as well as an orphan minor pilin, FimT, located elsewhere in the chromosome (Figure 1A). We therefore focused on interrogating the role of minor pilins in DNA binding in *A. baylyi*.

**Figure 1.**
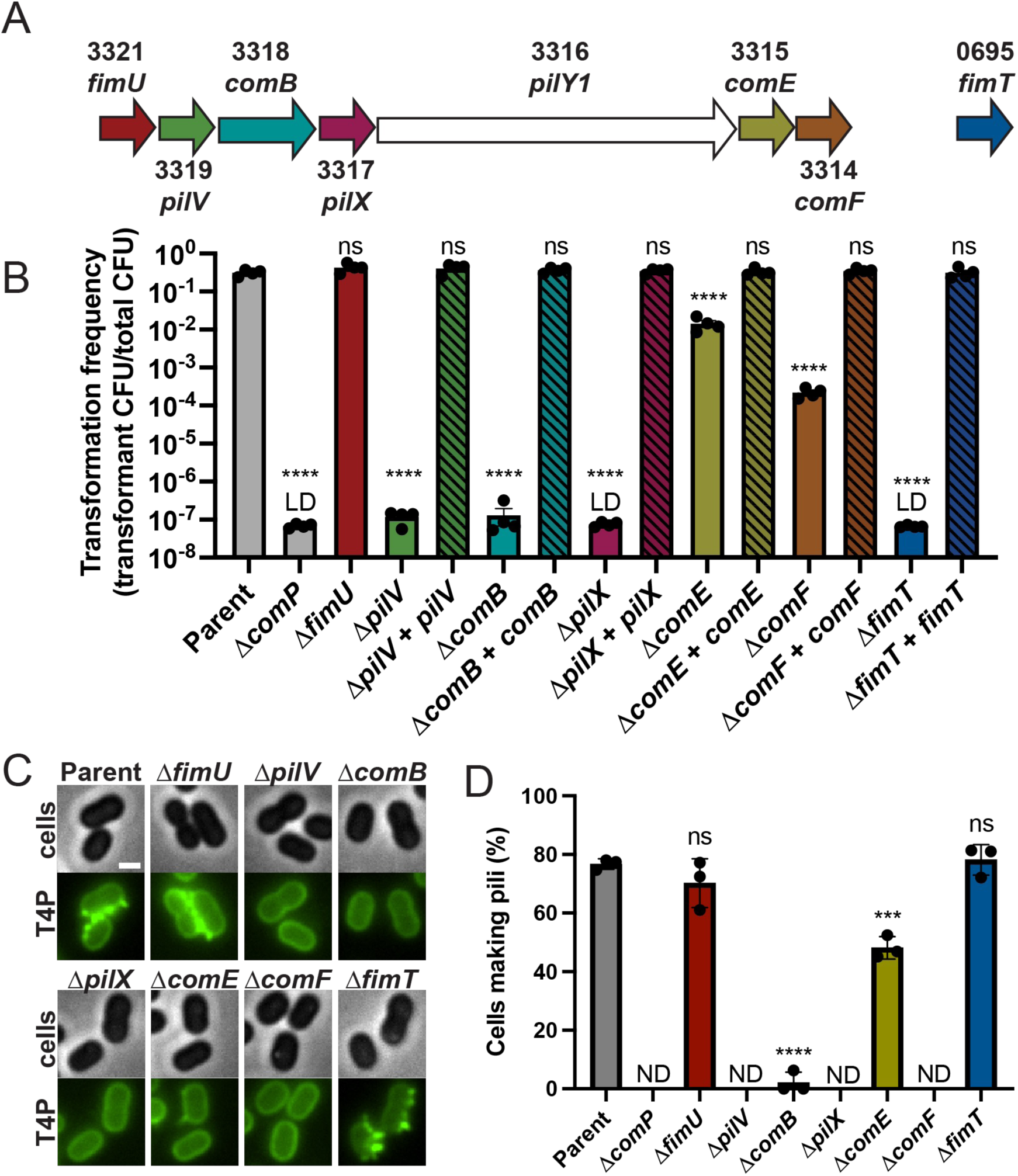
The minor pilin FimT is required for natural transformation but not for T4P synthesis. (A) Schematic of the minor pilin operon found in *A. baylyi*. Numbers indicate ACIAD locus ID for each gene. (B) Natural transformation assays of indicated strains. Each data point represents an independent, biological replicate and bar graphs indicate the mean ± SD. The transformation frequency of the Δ*comP, ΔpilX,* and *ΔfimT* strains was below the limit of detection, indicated by LD. Notably, one out of four replicates from the Δ*pilV* strain and three out of four replicates from the Δ*comB* strains fell below the limit of detection, LD. (C) Representative images of indicated strains with labeled T4P. Scale bar, 1 µm. (D) Quantification of the percentage of cells in populations of indicated strains making T4P. Each data point represents an independent, biological replicate and bar graphs indicate the mean ± SD. For each biological replicate, a minimum of 50 total cells were assessed. Statistics were determined by Dunnett’s multiple comparisons test to the parent strain. ****, *P* < 0.0001; ***, *P* < 0.001; ns, not significant.

## Results

To determine if specific minor pilins contribute to DNA binding, we created in-frame deletion mutants and assessed their ability to undergo natural transformation. All minor pilin mutant strains exhibited varying degrees of defects in transformability that could be fully complemented by ectopic expression constructs, with the exception of Δ*fimU* which has wildtype frequencies of natural transformation (Figure 1B). The high variability in transformation frequencies across different deletion strains indicates that different minor pilins may have divergent functions (Figure 1B). Because some minor pilins are essential for T4P synthesis, we next assessed T4P phenotypes in individual minor pilin mutants. To visualize T4P, we used a previously established T4P-imaging method that relies on the substitution of a cysteine residue in the major pilin subunit that can then be labeled by click-chemistry via fluorescent maleimide dyes (17, 18) (Figure 1C). Microscopy experiments showed that all minor pilin mutants except for Δ*fimU* and Δ*fimT* exhibit a defect in T4P synthesis, and we conclude that the deficit in natural transformation in *pilV*, *comB*, *pilX*, *comE*, and *comF* mutants can be explained by their reduction in T4P. Notably, we cannot exclude the possibility that these minor pilins may also contribute to DNA binding (Figure 1C, D). *fimU* mutants exhibited no defect in either natural transformation or T4P production, suggesting that FimU is involved in a different T4P function. *fimT* mutants exhibited no detectable transformation or defect in T4P synthesis, indicating that FimT may play a critical role in DNA binding.

Because most minor pilins are predicted to localize to the tips of T4P, we next assessed the localization of T4P-DNA interactions. Retraction motor mutants that are incapable of T4P retraction are hyperpiliated, so we used mutants lacking the retraction motor (Δ*pilT*) to improve our ability to detect T4P-DNA binding events. *A. baylyi* produces its T4P in a line along the long axis of the cell, limiting our ability to distinguish individual T4P filaments (19). To resolve individual T4P, we used a Δ*fimL* mutant background, which we previously showed has dispersed T4P localization but wildtype rates of natural transformation and T4P production (19) (Figure 2A). We fluorescently labeled DNA using commercially available dyes and evaluated DNA localization after incubation with hyperpiliated Δ*pilT* cells. For all instances where individual T4P could be resolved, DNA interactions occurred exclusively at the tips of T4P (Figure 2A). While minor pilins are generally predicted to localize to T4P tips, some studies have found that minor pilins can incorporate throughout the T4P fiber (20, 21). To assess the localization pattern of FimT, we performed immunofluorescence microscopy experiments with a strain encoding a largely functional (Figure S1) *fimT-3xFLAG* allele. As in fluorescent DNA experiments, FimT was found exclusively at the T4P tips in all instances where it colocalized with resolvable, individual T4P fibers (Figure 2B). These data are consistent with our previous results from *V. cholerae* where DNA binding occurs at the tips of T4P (6), supporting a broadly applicable model where T4P bind DNA via tip-localized minor pilins.

**Figure 2.**
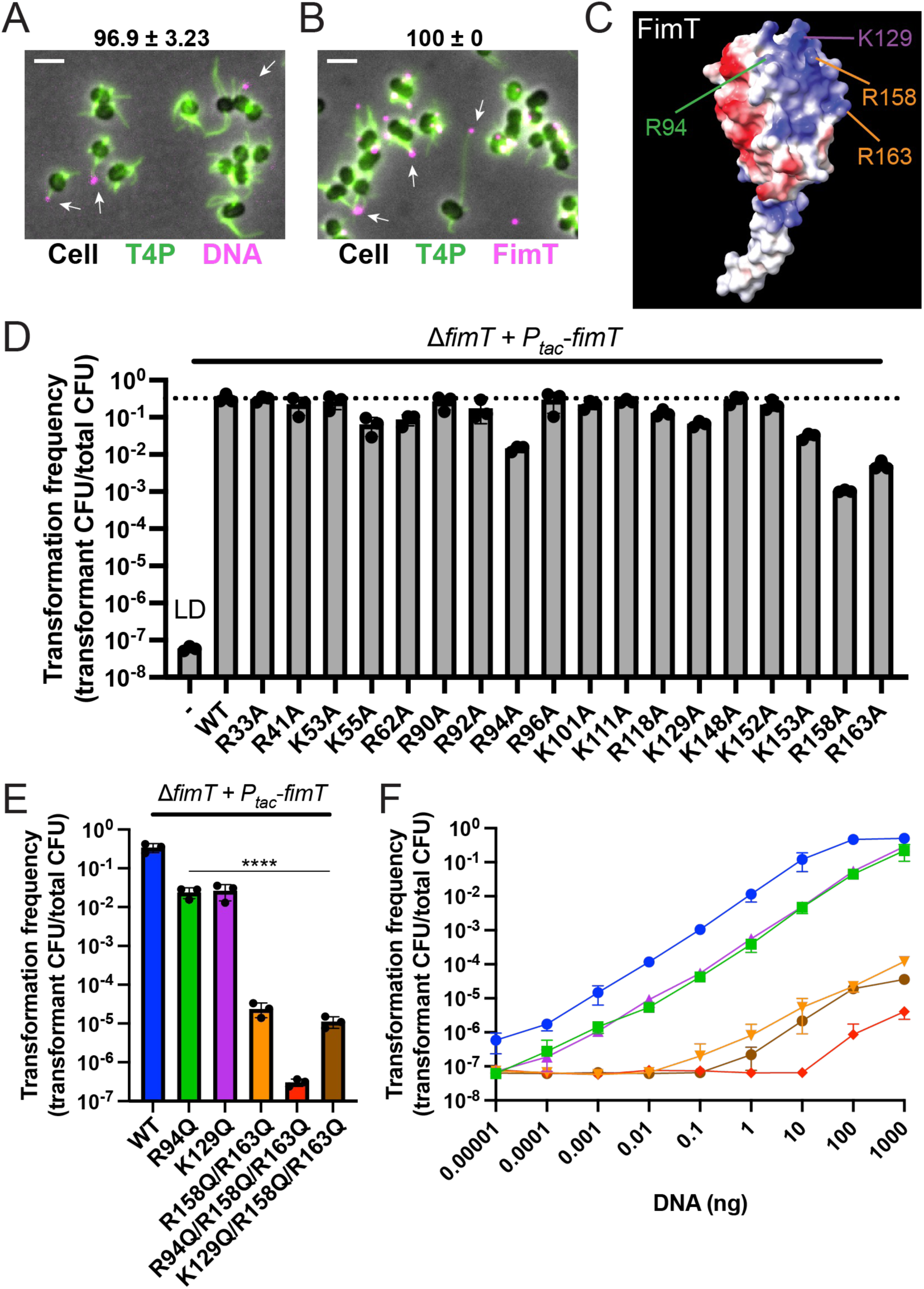
FimT binds DNA through electrostatic charges in *A. baylyi*. (A) Representative image of a Δ*fimL* Δ*pilT* strain with fluorescently labeled T4P (green) and DNA (fuchsia) showing DNA localization at T4P tips and (B) representative image of a Δ*fimL* Δ*pilT* strain with fluorescently labeled T4P (green) and immuno-labeled FimT (fuchsia) showing FimT localization at T4P tips. Scale bars, 2 μm. Numbers above panels indicate mean number of pili with fluorescent DNA or immunolabeled FimT at the tips ± SD of three biological replicates. A minimum of 30 pili with fluorescent puncta were quantified for each biological replicate. (C) AlphaFold-predicted structural model of FimT from A. baylyi showing regions of positively (blue) and negatively (red) charged regions. (D) Natural transformation assays of indicated strains. Each data point represents an independent, biological replicate and bar graphs indicate the mean ± SD. The transformation frequency of the uncomplemented *ΔfimT* strain was below the limit of detection, indicated by LD. (E) Natural transformation assays of indicated strains containing combinatorial mutations in FimT. Each data point represents an independent, biological replicate and bar graphs indicate the mean ± SD. Statistics were determined by Dunnett’s multiple comparisons test to the parent strain. ****, *P* < 0.0001. (F) DNA binding curve showing natural transformation results upon addition of different concentrations of DNA. Data show mean ± SD of three biological replicates.

We next speculated that FimT may bind DNA via electrostatic charges as it does in *V. cholerae* (6), and the AlphaFold structure of FimT predicts a large region of localized, positively charged residues that may interact with the negatively charged backbone of DNA (Figure 2C). We applied an unbiased genetic approach to identify positions that could mediate DNA binding by individually mutating all positively charged residues within FimT to uncharged residues. Transformation frequencies of FimT mutants show the importance of the C-terminal region of FimT in natural transformation in addition to other positively charged residues (Figure 2D). A recent study identified the C-terminus of a FimT homologue (sharing 27% identity and 57% similarity with *A. baylyi* FimT) as an important region for DNA binding during natural transformation in *Legionella pneumophila* which reaches transformation frequencies of ∼1 in 10,000 cells (22). In line with these results, our unbiased genetic approach revealed identical residues that play a considerable role in natural transformation in *A. baylyi* (prepilin positions R158/R163 corresponding to prepilin positions R153/R158 in the *L. pneumophila* FimT) (Figure S2A). However, in contrast to *L. pneumophila*, FimT from *A. baylyi* possesses additional positively charged residues (K55, K62, R94, K129) that appear to be important for natural transformation, and we predicted that these additional positively charged residues may allow for increased DNA binding.

To assess the role of different positively charged residues in DNA binding, we constructed strains containing combinatorial mutations in FimT, focusing on the two C-terminally encoded residues R158/R163 and the residues R94 or K129. Mutations to both R94 and K129 reduced transformation frequencies by 10-fold without affecting FimT stability (Figure 2E, S3). Combinatorial mutations in the C-terminal positively charged residues R158/R163 resulted in an additive decrease in transformation frequencies by 100-1000-fold lower than either individual mutant in contrast to *L. pneumophila* where mutations in both residues completely abrogated natural transformation (22) (Figure 2E). These data support a model where FimT from *A. baylyi* contains additional residues involved in DNA binding compared to other species. To determine the contribution of additional positively charged residues in *A. baylyi*’s FimT, we combined the R94Q and K129Q mutations with the C-terminal mutations R158Q and R163Q (Figure 2E). While the R94Q mutant exhibited reduced transformation almost to the limit of detection when combined with R158QR163Q, the addition of the K129Q mutation did not have a dramatic effect on natural transformation compared to the R158QR163Q mutant alone. These results suggest that there may be dedicated regions of positively charged residues within FimT that additively contribute to increased DNA binding. Where K129 acts in conjunction with C-terminal residues R158/R163 to coordinate stability of DNA binding in that area, the R94 residue may exist in an independent DNA binding pocket contributing to the overall increased DNA binding by FimT from *A. baylyi*.

Our transformation data indicate that FimT mutants possess different capacities for binding DNA, and so we performed DNA-titration transformation assays where we increased DNA concentrations to infer DNA binding by mutant alleles of FimT. In agreement with our hypothesis that our combinatorial mutants are defective in DNA binding, DNA titration curves show that increased levels of DNA consequently increase transformation rates across strains (Figure 2F). Increasing DNA concentration in 10-fold increments resulted in concomitant 10-fold increases in transformation frequencies across strains, with the strongest defect in DNA binding exhibited by the R94QR158QR163Q triple mutant. We next sought to directly quantify T4P-DNA binding in each mutant, and so we applied a microscopy approach to quantify fluorescently labeled DNA associated with cells. Microscopy experiments showed similar trends of DNA binding (Figure 3A) to transformation rates (Figure 2E), demonstrating that reduced DNA binding capability is responsible for the reduction in transformation frequencies in FimT mutants.

**Figure 3.**
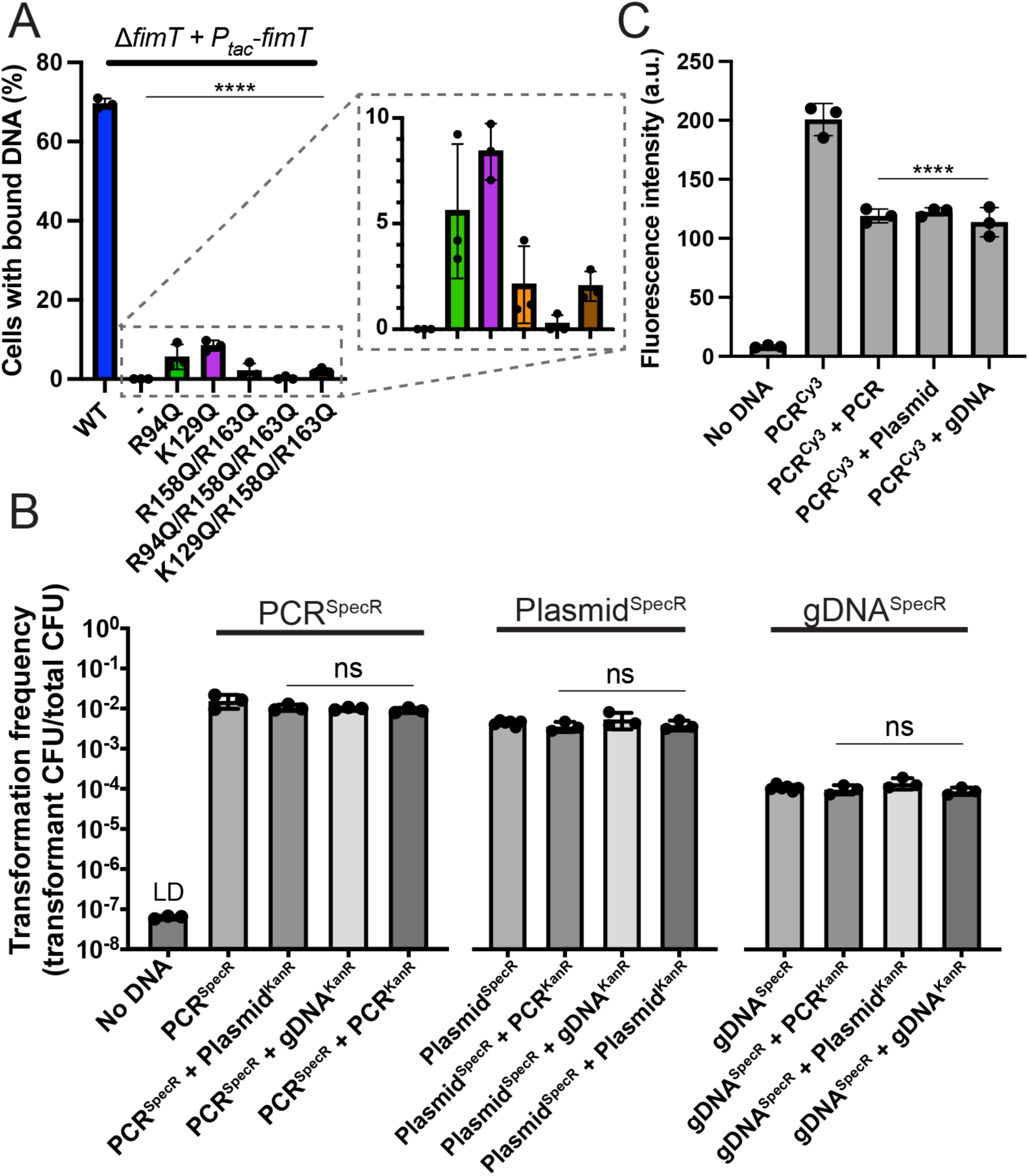
FimT binds nonspecific DNA. (A) DNA binding assay showing the proportion of cells associated with labeled fluorescent DNA in indicated strains. (B) Natural transformation assays performed with competing types of DNA in a 1:1 ratio reported as the transformation frequency of cells incorporating the transforming DNA containing the spectinomycin resistance cassette. Transformation frequencies for the competing transforming DNA containing the kanamycin resistance cassette from the same experiment can be found in supplemental figure 4. (C) Competition assays assessing DNA binding by fluorescently labeled PCR product competed against other unlabeled DNA types in a 1:1 ratio. Data points show mean ± SD of three independent, biological replicates. Statistics were determined by Dunnett’s multiple comparisons tests. For (A), DNA binding statistical comparisons were made to the Ptac-FimT^WT^ strain. For (B), PCR transformation comparisons were made to PCR^SpecR^, Plasmid transformation comparisons were made to Plasmid^SpecR^, and gDNA transformation comparisons were made to gDNA^SpecR^. For (C), statistical comparisons were made to PCR^Cy3^. ****, *P* < 0.0001; ns, not significant.

We reasoned that if electrostatic interactions mediate DNA binding, then DNA type (e.g. PCR product vs plasmid or genomic gDNA) should not influence FimT-DNA binding. To test this hypothesis, we performed transformation assays competing different types of DNA against each other in a 1:1 ratio by mass. We used different types of DNA carrying either spectinomycin or kanamycin resistance cassettes at the same genetic locus and determined transformation frequencies for both types of DNA (Figure 3B). To remove variability that could be introduced by transformations with replicating plasmids, we constructed a non-replicating plasmid carrying a homologous region equal in size to the transforming PCR products. We likewise used extracted gDNA carrying resistance markers at the same targeted locus to prevent variability that could arise from differences in the efficiency of recombination at different genetic loci. Under all competition conditions, we found that the type of DNA did not dramatically influence DNA uptake or transformability (Figure 3B, S4). To further test whether DNA type influences DNA binding, we used fluorescently labeled PCR product and competed it against non-fluorescent DNA of different types again in a 1:1 ratio. Non-fluorescent DNA regardless of type reduced fluorescent DNA signal by approximately half, as would be expected in a 1:1 fluorescent:non-fluorescent DNA mix (Figure 3C).

Since FimT from *A. baylyi* may bind DNA better than FimT homologues from other species, we wondered whether its expression could increase DNA binding and natural transformation in other bacteria. FimT is localized at the tips of T4P, suggesting that it forms a complex with other minor pilins that are also predicted to localize to T4P tips. Since proteins that interact co-evolve, we reasoned that FimT from *A. baylyi* (FimT^Ab^) would be unlikely to interact with minor pilins from distantly related species like *L. pneumophila*, making it difficult to interrogate function. We thus focused our efforts on closely related species with lower transformation frequencies to avoid issues that may arise from disruptions in protein-protein interactions in the T4P minor pilin complex. The clinical isolate *A. nosocomialis* strain M2 was originally classified as *Acinetobacter baumannii* (23) but exhibits less innate antibiotic resistance compared to more recent clinical isolates of *A. baumannii* (24). Both pathogens undergo natural transformation (12, 23), and their FimT homologues share 89% identity and 96% similarity (25) (Figure 4A, S2B) so we employed *A. nosocomialis* as a model system for ease of use. We first labeled T4P in *A. nosocomialis* using a functional pilin-cysteine knock-in strain (Figure S5) to test whether FimT was important for T4P synthesis in *Acinetobacter* pathogens. Similar to *A. baylyi*, a *fimT* deletion in *A. nosocomialis* did not impact T4P production (Figure 4B, C), but abolished natural transformation (Figure 4D). Ectopic complementation of its native FimT (FimT^An^) restored the defect in transformability to wildtype levels of ∼1 per 10,000 cells. In contrast, expression of FimT^Ab^ resulted in a 100-fold increase in transformation relative to the wildtype strain, reaching frequencies of ∼1 per 100 cells (Figure 4D). Collectively, these results demonstrate that DNA binding capacity of T4P sets a threshold for natural transformation.

**Figure 4.**
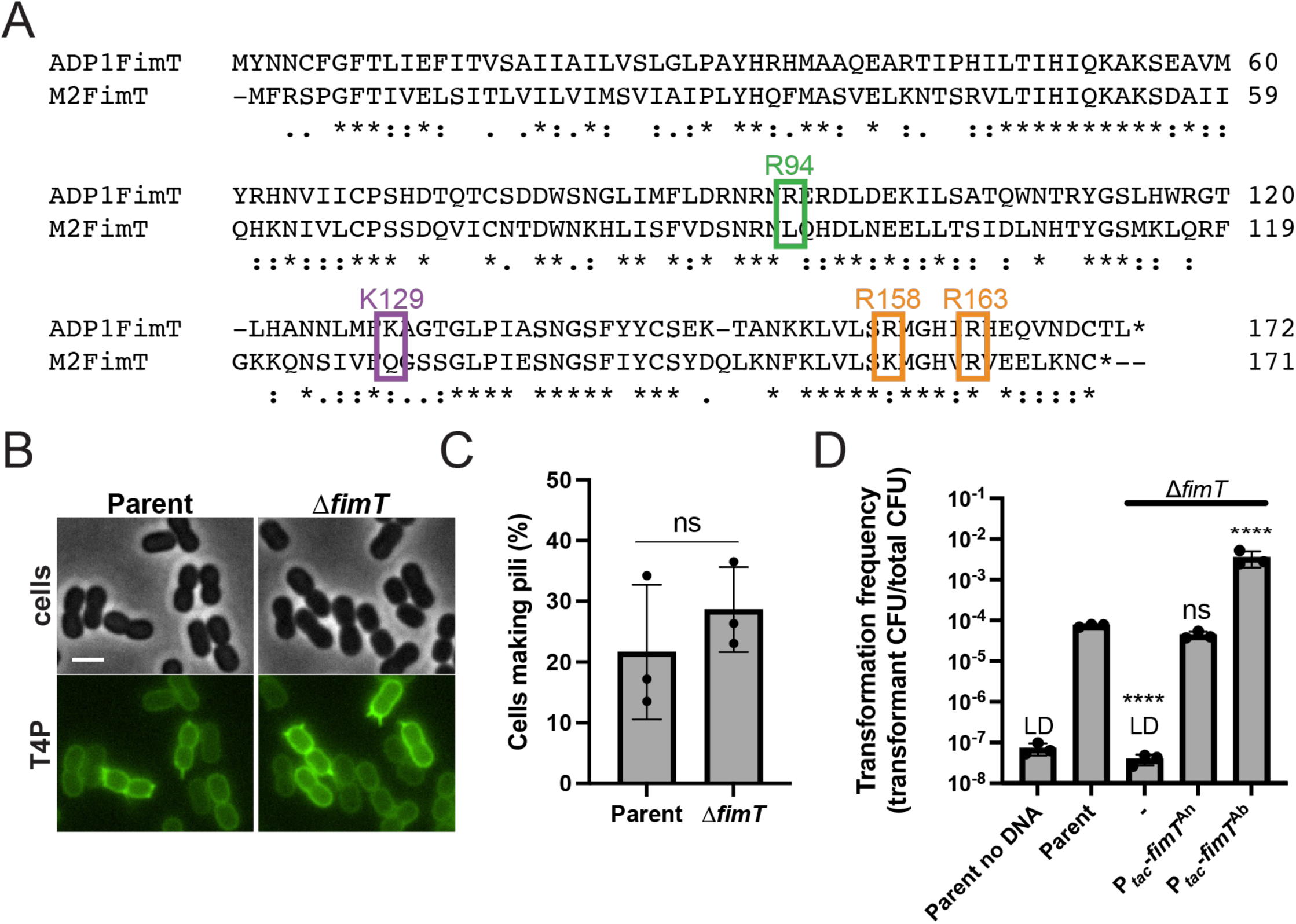
FimT from *A. baylyi* increases rates of natural transformation in the pathogen *Acinetobacter noscomialis*. (A) Alignment of *A. baylyi* strain ADP1 FimT (gene locus tag ACIAD0695) and *A. nosocomialis* strain M2 FimT (gene locus tag FDQ49_12665). Boxes indicate residues that are important for natural transformation in *A. baylyi.* ADP1FimT and M2FimT share 27% identity and 52% similarity. (B) Representative images of indicated strains with labeled T4P. Scale bar, 2 µm. (C) Quantification of the percentage of cells in populations of indicated strains making T4P. Each data point represents an independent, biological replicate and bar graphs indicate the mean ± SD. For each biological replicate, a minimum of 50 total cells were assessed. Statistics were determined between indicated strains by two-tailed t test. (D) Natural transformation assays of indicated strains. Each data point represents an independent, biological replicate and bar graphs indicate the mean ± SD. The transformation frequency of the no DNA control and *ΔfimT* strains was below the limit of detection, indicated by LD. Statistics were determined by Dunnett’s multiple comparisons test to the parent strain. ****, *P* < 0.0001; ns, not significant.

## Discussion

Many species that have not been reported to be naturally competent under laboratory settings carry competence genes, suggesting that the phylogenetic distribution of natural transformation may be significantly underestimated (1). There are several hypotheses for the fitness advantages that DNA uptake may confer including nutrient salvaging from environmental DNA, the acquisition of novel genes, and the maintenance of genome fidelity by homology matching with incoming DNA(26).

Although there are many fitness advantages that could be conferred through natural competency, there are also significant disadvantages as evidenced by the many defense systems bacteria carry to destroy foreign DNA. Our data provide insight into additional mechanisms restricting DNA uptake and horizontal gene transfer by limiting the efficiency of T4P-DNA binding. Many bacteria limit T4P-DNA binding by reducing T4P synthesis, but this may be disadvantageous under certain conditions since T4P are involved in multiple functions (5). Here we present a new mechanism whereby bacteria can alter T4P-DNA binding efficiency by incorporating a unique orphan pilin. This phenomenon could be imposed as a regulatory mechanism that provides an alternate pathway for regulating T4P-DNA binding while maintaining the production of T4P that can be used in other bacterial behaviors. We previously showed that *A. baylyi* also uses its T4P to interact with sister cells and control the spatial and structural development of three-dimensional multicellular communities (19). By employing a dedicated pilin for DNA uptake, *A. baylyi* could regulate DNA uptake capacity by limiting incorporation of FimT while still maintaining interactions with sister cells, highlighting a potential mechanism for how cells could control different T4P-related behaviors driven by the same T4P appendages.

There are several systems that can limit natural transformation downstream of DNA uptake. It was recently shown that some *Acinetobacter* pathogens possess restriction-modification (R-M) systems that recognize specific methylation patterns on incoming DNA to modulate transformation frequency (27). R-M systems found in *Helicobacter pylori* and *Campylobacter jujuni* likewise prevent transformation by incoming foreign DNA (28, 29). Intracellular nucleases targeting single-stranded DNA can also inhibit transformation rates (30) and CRISPR-Cas defense systems have been implicated in inhibiting natural transformation (31). The culmination of these studies highlights the importance of regulating DNA uptake, and here we identify a new mechanism for controlling natural competency through DNA binding efficiency.

In many species, *fimT* genes are found in orphan loci distant from other T4P minor pilin genes that are typically encoded together in a conserved operon (22) (Figure 1A). The genetic isolation of *fimT* indicates that it may have been horizontally acquired and that it could be transferred as a mobile T4P component via horizontal gene transfer to other species (32). Interestingly, other FimT homologues appear to lack the residues identified here that result in increased DNA binding, suggesting this characteristic increase in DNA binding by FimT might have evolved specifically in *A. baylyi*. Exploring the fitness advantages conferred by increased DNA binding in *A. baylyi* is the subject of future study. Our finding that different regions of FimT appear to coordinate DNA binding independently suggests that increasing the number of DNA binding sites in minor pilins from other species may permit increased DNA uptake, enabling the development of better genetic tools in genetically intractable model organisms.

Ultimately, these data establish DNA binding as a major limiting step in natural transformation, providing a foundation for the development of tools that can further hinder or boost DNA binding in other species.

## Materials and Methods

### Bacterial strains and culture conditions

*A. baylyi* strain ADP1 was used throughout this study and *A. nosocomialis* strain M2. For a list of strains used throughout, see Supplementary Table S1. *A. baylyi* cultures were grown at 30 °C in lysogeny broth (LB) medium (unless otherwise indicated) and on agar supplemented with kanamycin (50 µg/mL), spectinomycin (60 µg/mL), gentamycin (30 µg/mL), chloramphenicol (30 µg/mL), zeocin (100 µg/mL), and/or apramycin (50 µg/mL) as appropriate. *A. nosocomialis* cultures were grown at 37 °C in LB medium and on agar supplemented with kanamycin (50 µg/mL), gentamycin (30 µg/mL), and/or apramycin (50 µg/mL) as appropriate.

### Strain construction

#### Construction of mutant strains in *A. baylyi*

Mutants in *A. baylyi* were made using natural transformation as described previously (11, 19). Briefly, mutant constructs were made by splicing-by-overlap (SOE) PCR to stich (1) ∼3 kb of the homologous region upstream of the gene of interest, (2) the mutation where appropriate (for example deletion by allelic replacement with an antibiotic [AbR] cassette), and (3) ∼3 kb of the homologous downstream region. For a list of primers used to generate mutants in this study, see Supplementary Table S2. The upstream region was amplified using F1 + R1 primers, and the downstream region was amplified using F2 + R2 primers. All AbR cassettes were amplified with ABD123 (ATTCCGGGGATCCGTCGAC) and ABD124 (TGTAGGCTGGAGCTGCTTC). Fusion proteins were amplified using the primers indicated in Supplementary Table S2. In-frame deletions were constructed using F1 + R1 primer pairs to amplify the upstream region and F2 + R2 primer pairs to amplify the downstream region with ∼20 bp homology to the remaining region of the downstream region built into the R1 primer and ∼20 bp homology to the upstream region built into the F2 primer. SOE PCR reactions were performed using a mixture of the upstream and downstream regions, and middle region where appropriate using F1 + R2 primers. SOE PCR products were added with 50 µL of overnight-grown culture to 450 µL of LB in 2-mL round-bottom microcentrifuge tubes (USA Scientific) and grown at 30 °C rotating on a roller drum for 3–5 h. For AbR-constructs, transformants were serially diluted and plated on LB and LB + antibiotic. For in-frame deletions and protein fusion constructs, after the 3-5 h incubation, cells were diluted and 100 µL of 10^-6^ dilution was plated on LB plates. In-frame deletions were confirmed by PCR using primers ∼150 bp up- and downstream of the introduced mutation, and fusions were confirmed by sequencing. For construction of mutants containing multiple mutations that each result in transformation frequencies below the limit of detection (for example Δ*fimT* Δ*pilT* mutants), the parent strain was co-transformed simultaneously with PCR products containing both mutations as described above and selected for on appropriate antibiotic plates. Each mutation introduced by co-transformations was confirmed appropriately as described above.

Complementation strains were constructed by placing the gene of interest under a constitutive P^tac^ promoter at the *vanAB* locus as described previously (11). First, the *vanAB* locus containing the KanR and the constitutive P*^tac^* promoter published previously (11) was amplified using primers CE173 + CE336 to amplify the upstream region. The downstream region was amplified from the same KanR-P*^tac^* construct strain using primers CE406 + CE176. The complementation gene was then amplified using specified primers found in Supplementary Table S2. SOE PCRs and natural transformations were then performed exactly as described above. To generate *fimT* point mutations, we first constructed a P*^tac^*-*fimT* strain at the *vanAB* locus as described above. We then used primers amplifying the up and downstream regions including the P*^tac^*-*fimT* region with point mutations built into the R1 and F2 primers and carried out transformations and sequencing as described above. Because most minor pilin mutants including Δ*fimT* are not transformable, we first constructed *vanAB* expression constructs and then transformed the deletion into the expression construct strain and confirmed the deletion region by PCR and the *vanAB* locus by sequencing.

To make the P*^tac^*-*fimT*-*3xFLAG* strain, we used genomic material from our P*^tac^*-*fimT* strain as a DNA template, amplifying the upstream region including *fimT* without the stop codon with primers CE173 + CE2210 and the downstream region using primers CE2135 + CE176 where the linker for the 3xFLAG tag was built into CE2210 and CE2135. We then performed a second round of PCRs to introduce the 3xFLAG tag to our DNA by re-amplifying the upstream region with primers CE173 + CE1778 and the downstream region with primers CE1777 + CE176 with the 3xFLAG tags built into primers CE1777 and CE1778. This strain was confirmed by sequencing, and then its genomic material was used as a template to make the point mutant derivatives as described above.

#### Construction of mutant strains in *A. nosocomialis*

Mutants in *A. nosocomialis* were made using natural transformation and SOE products similarly to *A. baylyi* as described above. However, *A. nosocomialis* is not transformable in liquid media and instead requires a surface to induce competency. Strains were grown by shaking overnight in LB broth at 37 °C. Then, 50 µL of overnight culture was subcultured into 3 ml of LB broth and grown shaking for 2-3 h until cells were in exponential growth phase. 1 µL of exponential culture was diluted into 100 µL of phosphate-buffered saline (PBS), and then 5-10 µL of diluted culture was mixed with ∼50 ng of transforming DNA and spotted onto the surface of transformation agar (TA) (2.5 g/L NaCl, 5 g/L tryptone, 2% agarose) in microcentrifuge tubes and incubated overnight at 30 °C. The next day, cells were removed from the surface of the TA using 200 µL of PBS to resuspend surface-associated cells. Cells were then diluted and plated on LB + antibiotic plates and grown overnight at 37 °C.

Because natural transformation rates in *A. nosocomialis* are significantly lower than those of *A. baylyi*, we inserted an apramycin or kanamycin resistance cassette upstream of the *pilA* locus to generate cysteine point mutations. To generate this *pilA*::*AbR*, *pilA* strain, we used primers CE703 (F1) + CE944 (R1) to amplify the upstream DNA and primers CE943 (F2) + CE744 (R2) to amplify the downstream DNA from the wildtype strain. We amplified an resistance cassettes using primers ABD123 + ABD124 and then transformed the SOE product into *A. nosocomialis* and selected for the *pilA*::*AbR*, *pilA* strain on appropriate plates. The strains carrying the AbR cassettes upstream of *pilA* was confirmed by sequencing and then used as a template for generating SOE PCRs carrying point mutations in *pilA* built into the R1 and F2 primers the same way as described above for *A. baylyi* and detailed in Supplemental Table S2. Complementation constructs in *A. nosocomialis* were also generated at the *vanAB* locus as in *A. baylyi*. P*^tac^* expression constructs were first built into *A. baylyi* at the *vanAB* locus as described above and then the DNA carrying the antibiotic resistance cassette and the P*^tac^* promoter were amplified using primers ABD123 + ABD124. The P*^tac^*-gene PCR product was then inserted into the *vanAB* locus DNA via SOE PCR, where the upstream DNA was generated using primers CE1727(F1) + CE1728 (R1) and the downstream DNA was generated using primers CE1729 (F2) + CE688 (R2). SOE products were then transformed into *A. nosocomialis*, selected for on kanamycin, and confirmed by sequencing.

#### Construction of plasmids

The plasmid pXB300 (33) that is nonreplicating in *A. baylyi* was used to construct transforming plasmids used in DNA competition assays. First, pXB300 was amplified using primers CE1449 + CE1450 and the insert regions containing ACIAD1551::AbR were amplified from strains carrying either kanamycin or spectinomycin cassettes using primers CE1467 + CE1468. Then, amplified plasmid and insert PCR products were assembled using NEBuilder HiFi DNA Master Mix (New England Biolabs) following manufacturer’s protocols and transformed into *Escherchia coli* strain S-17 by electroporation for maintenance and replication. *E. coli* was grown in LB medium supplemented with kanamycin (50 µg/mL) or spectinomycin (60 µg/mL) where appropriate at 37 °C.

### Natural transformation assays

Transformation assays in *A. baylyi* were performed exactly as previously described (11, 19). Briefly, strains were grown overnight in LB broth at 30 °C on a roller drum. Then, 50 µL of overnight culture was subcultured into 450 µL of fresh LB medium and at least 50 ng of transforming DNA (tDNA) (a ∼7 kb PCR product containing Δ*pilT*::*spec* amplified using primers CE49 + CE50) was used for all transformation assays except for DNA competition assays which are detailed below. DNA was quantified using a Qubit (ThermoFisher) following standard Qubit protocols.

Reactions were incubated with end-over-end rotation on a roller drum at 30 °C for 5 h and then plated for quantitative culture on LB + antibiotic plates (to quantify transformants) and on plain LB plates (to quantify total viable counts). Data are reported as the transformation frequency, which is defined as the (CFU/mL of transformants) / (CFU/mL of total viable counts).

Transformation assays in *A. nosocomialis* were performed on TA exactly as described above under mutant construction and quantified the same way as *A. baylyi* to determine transformation frequencies. The transforming DNA used for transformations in *A. nosocomialis* was produced by PCR by amplifying a ∼7 kb region containing a gentamycin resistance cassette at locus tag FDQ49_10705 which encodes a non-functional frame-shifted gene using primers CE1788 + CE1791.

### Pilin labeling, imaging, and quantification

Pilin labeling in *A. baylyi* was performed as described previously (19). Briefly, 100 µL of overnight cultures was added to 900 µL of fresh LB in a 1.5 mL microcentrifuge tube, and cells were grown at 30 °C rotating on a roller drum for 70 min. Cells were then centrifuged at 18,000 x *g* for 1 min and resuspended in 50 µL of LB before labeling with 25 µg/mL of AlexaFluor488 C5-maleimide (AF488-mal) (ThermoFisher) for 15 min at room temperature. Labeled cells were centrifuged, washed three times with 100 µL of PBS and resuspended in 5–20 µL PBS. Cell bodies were imaged using phase-contrast microscopy while labeled pili were imaged using fluorescence microscopy on a Nikon Ti2-E microscope using a Plan Apo 100X oil immersion objective, a GFP/FITC/Cy2 filter set for pili, a Hamamatsu ORCA-Fusion Gen-III cCMOS camera, and Nikon NIS Elements Imaging Software. Cell numbers and the percent of cells making pili were quantified manually using Fiji. All imaging was performed under 1% agarose pads made with PBS solution.

For labeling pili in *A. nosocomialis*, cells were treated as described above with a few differences. 50 µL of overnight cultures were diluted into 3 mL of LB and grown for 2-3 h to exponential growth phase. 1 mL of liquid exponential culture was then centrifuged and cells were labeled and imaged exactly as described above.

### Immunofluorescence and DNA localization in Δ*fimL* Δ*pilT* mutants

We previously found that overnight cultures of *A. baylyi* grown in M63 minimal medium + casamino acids and glucose (M63CA+G) (2 g/L (NH^4^)^2^SO^4^, 13.6 g/L KH^2^PO^4^, 0.5 mg/L FeSO^4^, 1 mM MgSO^4^, 0.5% w/v casamino acids, and 0.4% w/v glucose) produce longer pili than cells grown in LB (11). We therefore grew cells in M63CA+G medium for assessing DNA and FimT localization on *A. baylyi* pili as this increase in length vastly improved our ability to resolve individual pilus fibers. For DNA localization experiments, 100 µL of overnight cultures grown in M63CA+G were co-incubated with 25 µg/mL of AF488-mal and 100 ng of fluorescently labeled DNA for 30 min. Fluorescent DNA^Cy3^ was made from a ∼7 kb Δ*pilT*::*spec* PCR product that was fluorescently labeled as described previously (6) using the Cy3 LabelIT kit (Mirus Biosciences) as per the manufacturer’s recommendations. Cells were washed 3X with 100 µL of PBS before final resuspension in 20 µL PBS before imaging. For immunofluorescence experiments, 100 µL of overnight cultures grown in M63CA+G was centrifuged and resuspended in 100 µL of PBST (PBS + 0.05% v/v Tween20) containing a 1:100 dilution of mouse monoclonal α-FLAG antibodies (Sigma) and incubated overnight at 4 °C with end-over-end rotation in a microcentrifuge tube. The next day, cells were washed 3X with 200 µL cold PBST before resuspension in 100 µL cold PBST containing a 1:200 dilution of goat α-mouse antibodies conjugated to AlexaFluor555 (Abcam). Cells were incubated for 1 h at 4 °C with end-over-end rotation before washing 3X with 200 µL cold PBST. Cells were then resuspended into 50 µL cold PBST and incubated with 25 µg/mL of AF488-mal for 15 min. Cells were then washed 1X with plain PBS (Tween20 was omitted here as it has considerable background fluorescence) and then resuspended in 5-20 µL of plain PBS and imaged.1 µl of cells was used for microscopy using the same microscopy setup described above, using a DsRed/TRITC/Cy3 filter set (Nikon) to image both DNA^Cy3^ and FimT.

### Western blotting

Approximately 10^9^ cells from overnight cultures were concentrated into a pellet by centrifugation, and the culture supernatant was removed. Cell pellets were resuspended in 75 µL PBS and then mixed with 25 µL of 4X SDS-PAGE sample buffer (250 mM Tris, pH 6.8, 40% glycerol, 8% SDS, 0.8% bromophenol blue, and 20% β-mercaptoethanol) and boiled using a thermal cycler set to 99 °C for 10 min. Proteins were separated on a 4–20% pre-cast polyacrylamide gel (Biorad) by SDS electrophoresis, electrophoretically transferred to a nitrocellulose membrane, and probed with 1:5000 dilution of mouse monoclonal α-FLAG antibodies (Sigma) and a 1:12,000 dilution of mouse monoclonal α-RpoA (BioLegend) primary antibodies. Blots were washed and then probed with a 1:10,000 dilution of goat α-mouse antibody conjugated to horse radish peroxidase (Sigma). Blots were then incubated with SuperSignal West Pico PLUS Chemiluminescence substrate (ThermoFisher) and then imaged using a BioRad Chemidoc imaging system.

### Fluorescent DNA binding assays

100 µL of overnight-grown cultures was added to 3mL of fresh LB in a 14 mL culture tube (Corning) and cells were grown at 30 °C rotating on a roller drum for 2–3 h. Cells were then centrifuged at 18,000 x *g* for 1 min and resuspended in 50 µL of PBS, and 100 ng of DNA^Cy3^ was added and the mixture was incubated at room temperature for 30 min. To visualize DNA binding in *pilT* mutants, cells were grown as described above, but 100 ng of DNA^Cy3^ was added to cells along with AF488-mal and incubated for 15–25 min before washing. Cells were centrifuged at 18,000 x *g* for 1 min and washed three times with 200 µl of PBS and resuspended in 200 µL of PBS. 1 µl of cells was used for microscopy using the same microscopy setup described above, using a DsRed/TRITC/Cy3 filter set (Nikon) to image DNA^Cy3^ and the remaining sample was used to acquire fluorescence readings on a Synergy H1 multimode reader (BioTek) using Gen 6 software. For DNA type binding competition assays, 100 ng of DNA^Cy3^ was added to the parent strain along with 100 ng of unlabeled ∼7 kb PCR product, plasmid, or gDNA (a 1:1 ratio) and incubated for 30 minutes. Samples were then washed the same as above and then measured for fluorescence intensity on the Synergy H1 multimode reader.

### DNA competition assays

DNA competition assays were performed exactly as transformation assays described above with some differences. Briefly, strains were grown overnight in LB broth at 30 °C on a roller drum. Then, 50 µL of overnight culture was subcultured into 450 µL of fresh LB medium containing 5 ng of DNA of different types (for example 5 ng of PCR product and 5 ng of plasmid) carrying either a kanamycin or spectinomycin resistance cassette targeting the ACIAD1551 locus encoding a non-functional frameshifted gene. PCR product tDNA consisted of a ∼5 kb region amplified using primers CE1467 + CE1468 that were also used to construct the pXB300-1551::AbR plasmids. Plasmids were extracted using a standard Qiagen Miniprep kit and genomic DNA (gDNA) was extracted using a standard Wizard DNA Extraction Kit (Promega) and all DNA was quantified using a Qubit. All other aspects of this assay were performed exactly the same as transformation assays described above.

### Reproducibility

All experiments were repeated a minimum of two times, and all attempts at replication were successful.

## Acknowledgements

We are indescribably grateful for resources and laboratory space provided by M. S. Trent during early work on this project. We would like to thank J. Engel for providing the *A. nosocomialis* M2 strain, and we would like to thank X. Charpentier for providing the apramycin resistance gene and helpful tips for work with *Acinetobacter* pathogens. We would also like to thank A. B. Dalia and K. R. Hummels for critical comments on the manuscript. Courtney Ellison, PhD, is a Damon Runyon-Marilyn and Scott Urdang Breakthrough Scientist supported by the Damon Runyon Cancer Research Foundation (DFS6023). This work was supported by National Institutes of Health grant R35GM150916 awarded to C.K.E.

## Author contributions

C.K.E. designed and coordinated the overall study. C.K.E. and T.J.E. performed the experiments. C.K.E. and T.J.E. analyzed and interpreted data. C.K.E. wrote the manuscript with input from T.J.E.

## Data availability statement

Source data are provided in this paper.

---

**Figure S1.**
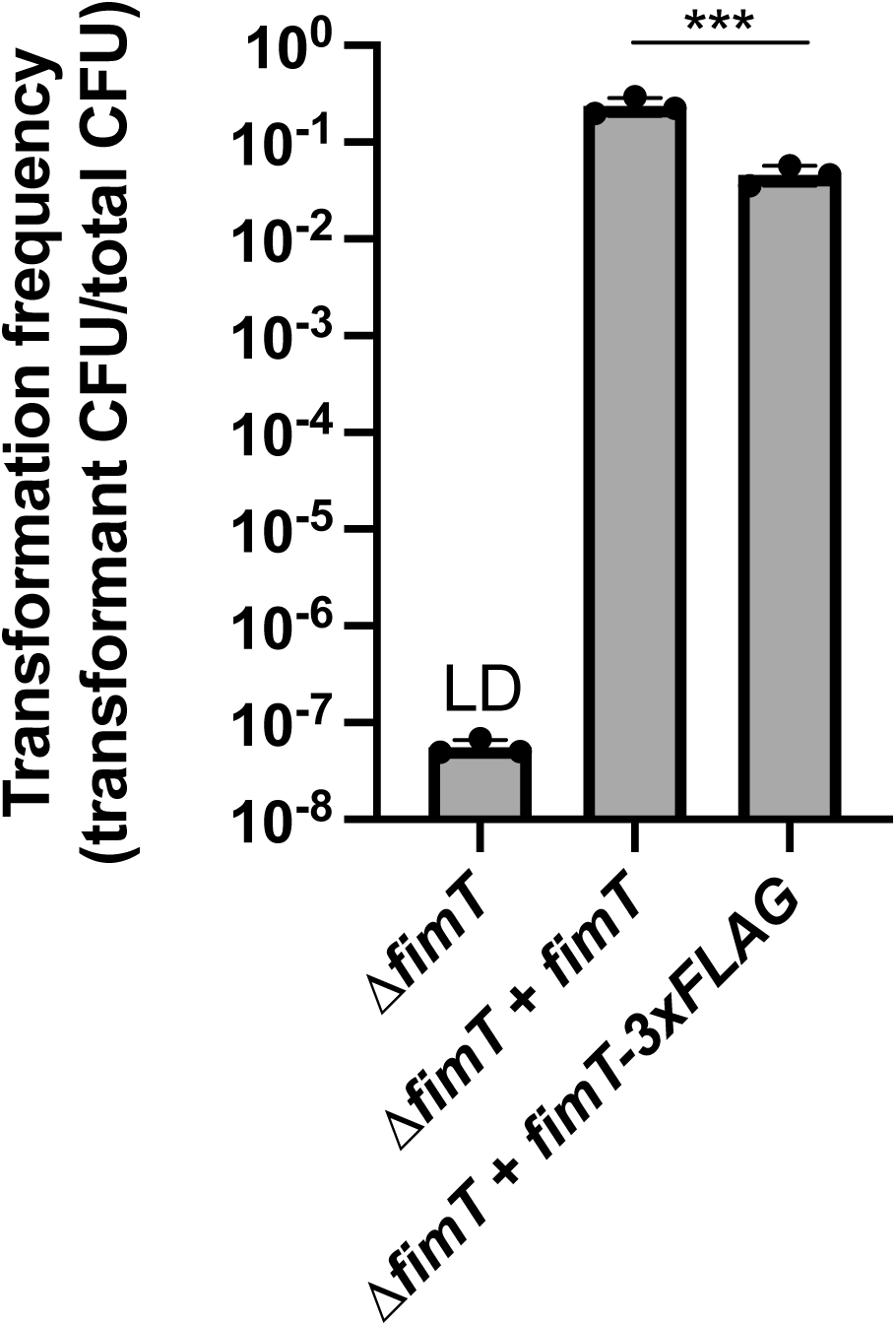
FimT-3xFLAG is largely functional. Natural transformation assays of indicated strains. Each data point represents an independent, biological replicate and bar graphs indicate the mean ± SD. The transformation frequency of the *ΔfimT* strain was below the limit of detection, indicated by LD. Statistics were determined between indicated strains by two-tailed t test of the log transformed data. ***, *P* < 0.001.

**Figure S2.**
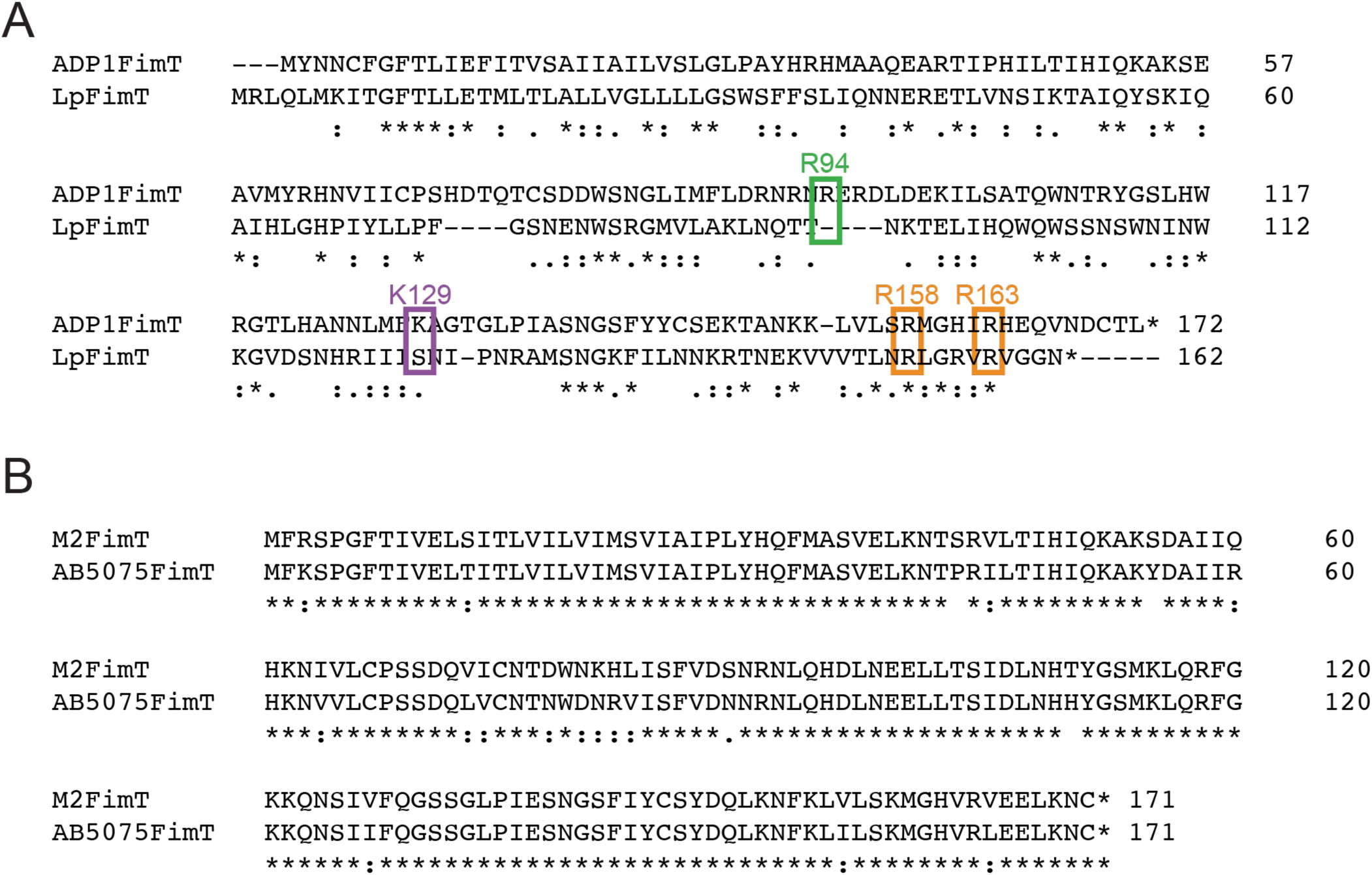
ClustalW alignments of prepilin protein sequences of FimT homologues. (A) Alignment of *A. baylyi* FimT (gene locus tag ACIAD0695) and *L. pneumophila* FimT (gene locus tag Lpg1428). Boxes indicate residues that are important for natural transformation in *A. baylyi.* LpFimT and ADP1FimT share 27% identity and 52% similarity. (B) Alignment of *Acinetobacter nosocomialis* M2 FimT (gene locus tag FDQ49_12665) with *Acinetobacter baumannii* AB5075 FimT (gene locus tag ABUW_0648). M2FimT and AB5075FimT share 89% identity and 96% similarity.

**Figure S3.**
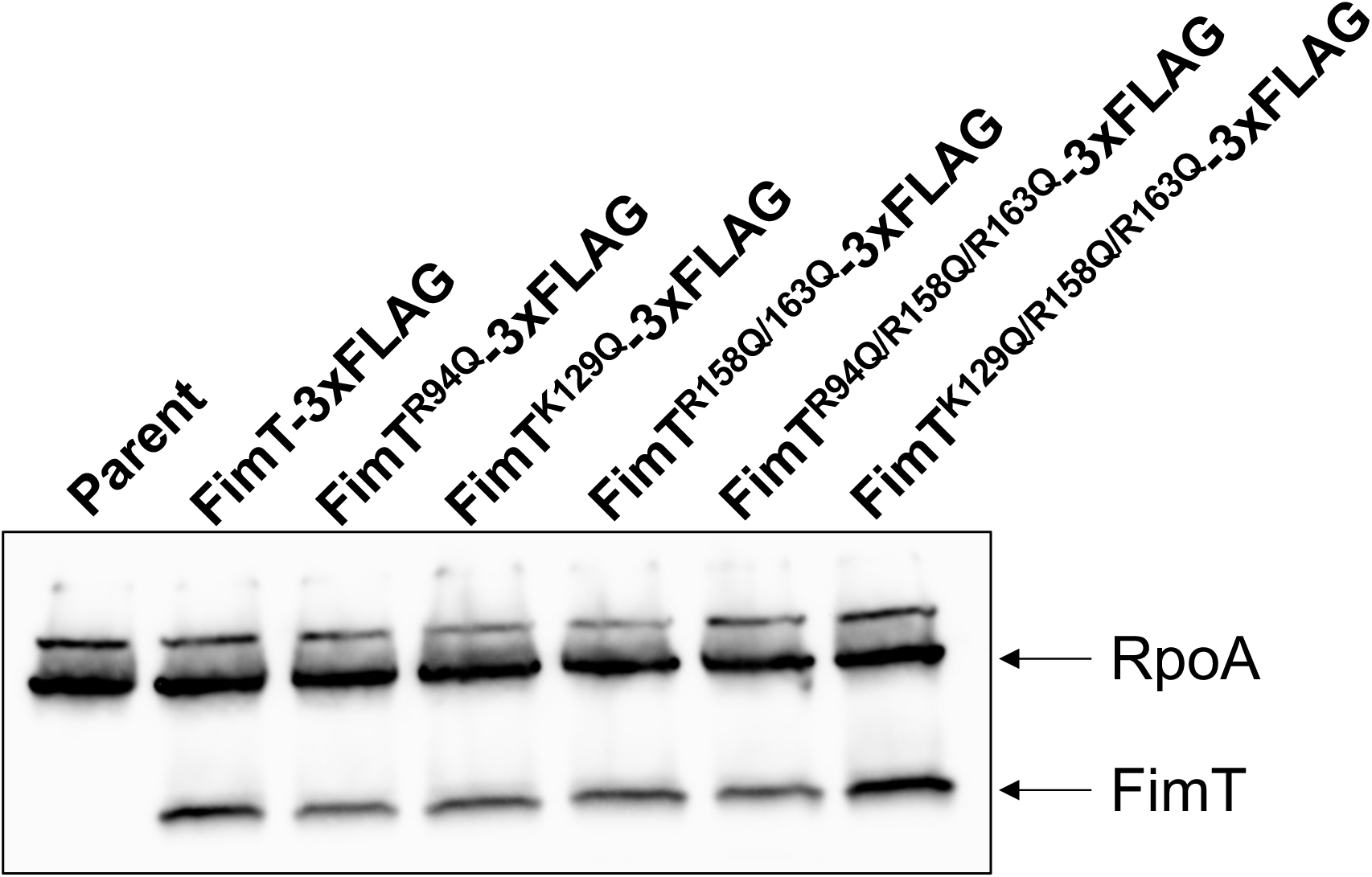
FimT point mutations affecting natural transformation do not affect stability or production of FimT. Point mutations were introduced to FimT-3xFLAG for Western blot analysis using α-FLAG antibody. α-RpoA antibody was used as a protein loading control.

**Figure S4.**
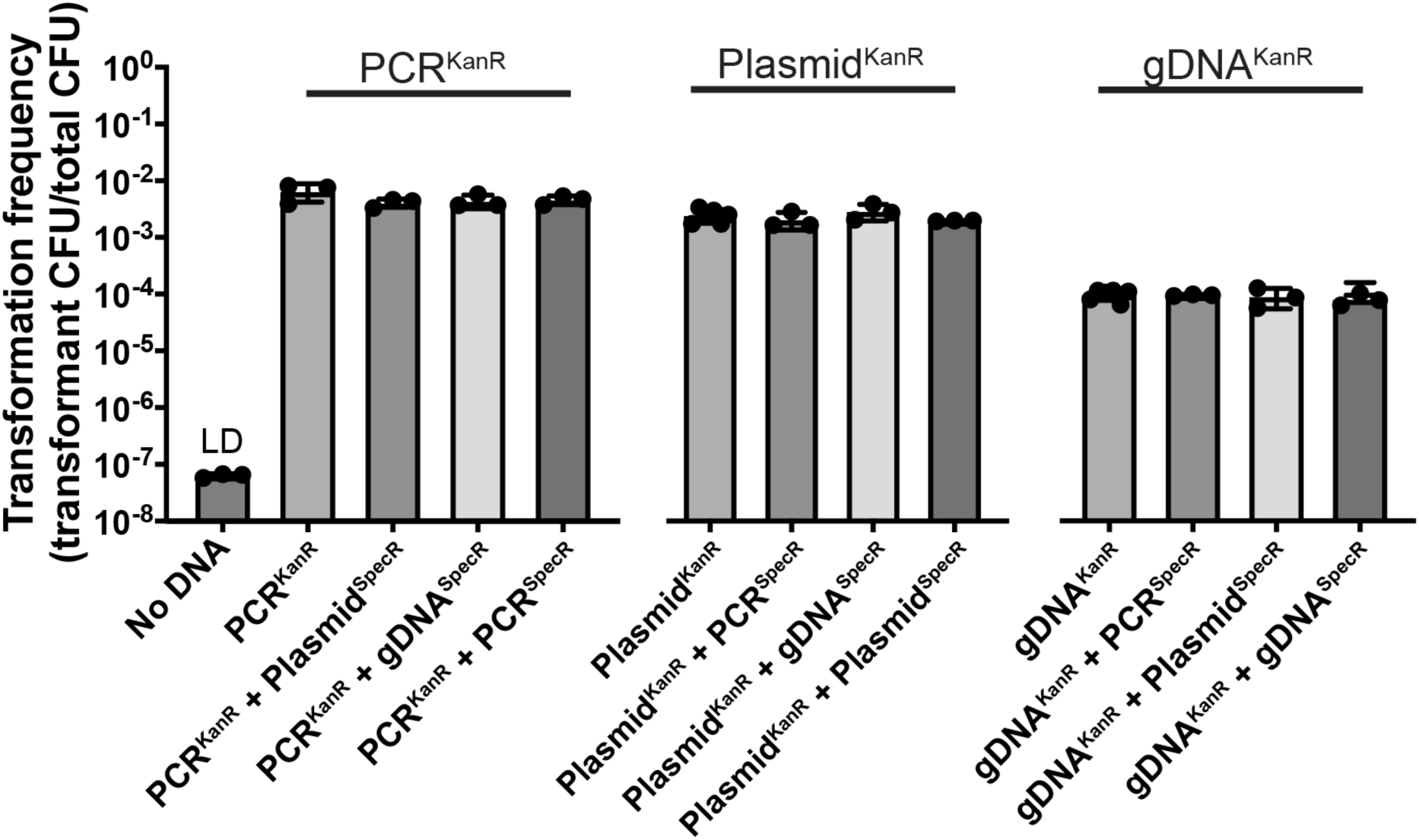
FimT binds nonspecific DNA. Natural transformation assays performed with competing types of DNA in a 1:1 ratio reported as the transformation frequency of cells incorporating the transforming DNA containing the kanamycin resistance cassette. Transformation frequencies for the competing transforming DNA containing the spectinomycin resistance cassette from the same experiment can be found in Figure 3B. Data points show mean ± SD of three independent, biological replicates.

**Figure S5.**
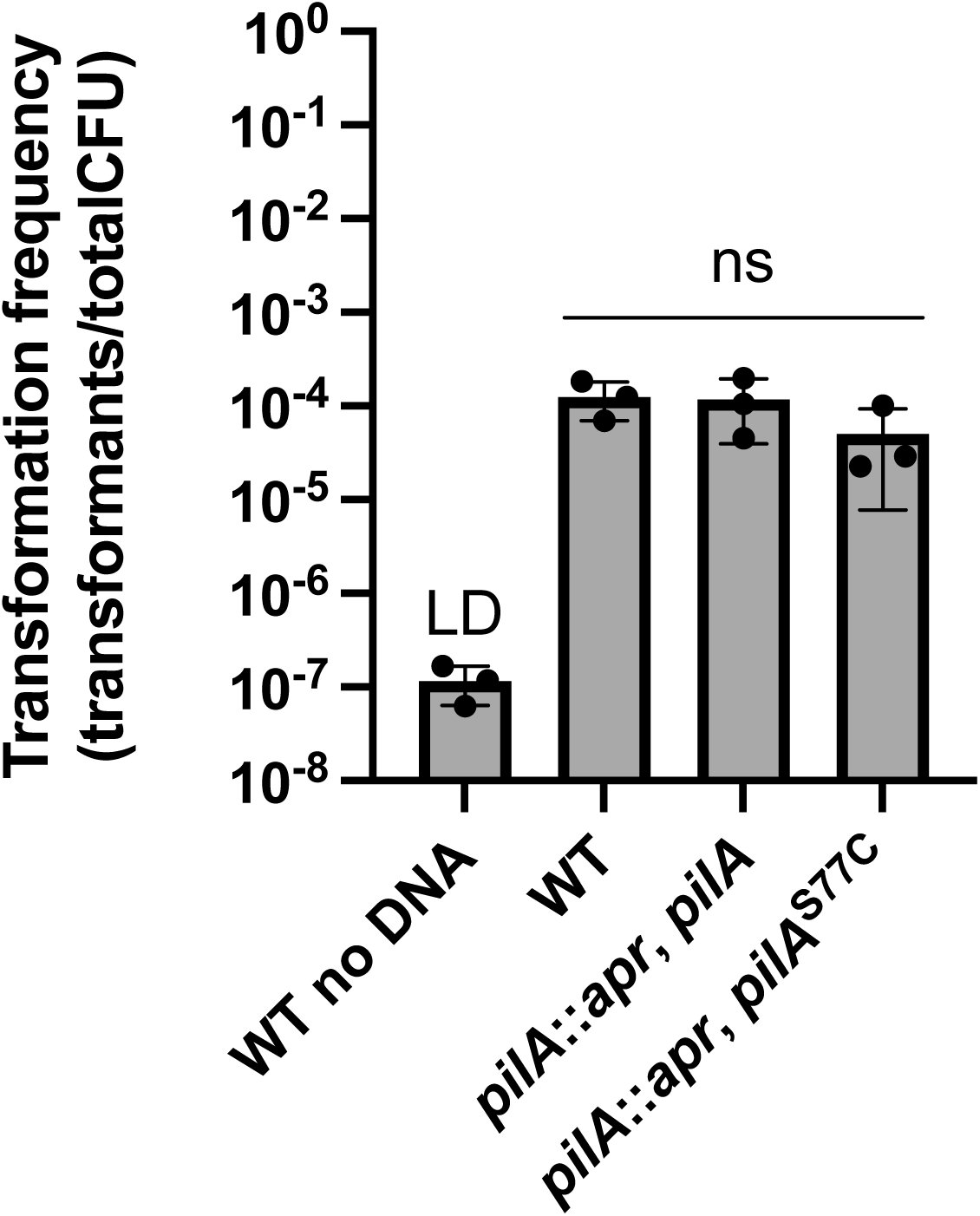
The *Acinetobacter nosocomialis* M2 pil-cys strain is functional. Natural transformation assays of indicated strains. Each data point represents an independent, biological replicate and bar graphs indicate the mean ± SD. The transformation frequency of the WT strain incubated without DNA exhibited no detectable colonies, indicated by LD. Statistics were determined by Dunnett’s multiple comparisons test to the WT strain. ns, not significant.

**Table S1.** Bacterial strains used in this study.

| Strain # | Strain name in manuscript | Genotype | Figures where strain appears | Ref |
| --- | --- | --- | --- | --- |
| CE100 | Parent | *A. baylyi* ADP1 *comP*<sup>T129C</sup> | 1B, 1C, 1D, 3B, S4 | 1 |
| CE358 | $\Delta comP$ | ADP1 $\Delta comP::kan$ | 1B, 1D | 1 |
| CKE1 | $\Delta fimU$ | ADP1 *comP*<sup>T129C</sup> $\Delta fimU$ | 1B, 1C, 1D | This study |
| CKE2 | $\Delta pilV$ | ADP1 *comP*<sup>T129C</sup> $\Delta pilV$ | 1B, 1C, 1D | This study |
| CKE3 | $\Delta pilV + pilV$ | ADP1 *comP*<sup>T129C</sup> $\Delta pilV \Delta vanAB::kan$, *P*<sub>tac</sub>-*pilV* | 1B | This study |
| CKE4 | $\Delta comB$ | ADP1 *comP*<sup>T129C</sup> $\Delta comB$ | 1B, 1C, 1D | This study |
| CKE5 | $\Delta comB + comB$ | ADP1 *comP*<sup>T129C</sup> $\Delta comB \Delta vanAB::kan$, *P*<sub>tac</sub>-*comB* | 1B | This study |
| CKE6 | $\Delta pilX$ | ADP1 *comP*<sup>T129C</sup> $\Delta pilX$ | 1B, 1C, 1D | This study |
| CKE7 | $\Delta pilX + pilX$ | ADP1 *comP*<sup>T129C</sup> $\Delta pilX \Delta vanAB::kan$, *P*<sub>tac</sub>-*pilX* | 1B | This study |
| CKE8 | $\Delta comE$ | ADP1 *comP*<sup>T129C</sup> $\Delta comE$ | 1B, 1C, 1D | This study |
| CKE9 | $\Delta comE + comE$ | ADP1 *comP*<sup>T129C</sup> $\Delta comE \Delta vanAB::kan$, *P*<sub>tac</sub>-*comE* | 1B | This study |
| CKE10 | $\Delta comF$ | ADP1 *comP*<sup>T129C</sup> $\Delta comF$ | 1B, 1C, 1D | This study |
| CKE11 | $\Delta comF + comF$ | ADP1 *comP*<sup>T129C</sup> $\Delta comF \Delta vanAB::kan$, *P*<sub>tac</sub>-*comF* | 1B | This study |
| CKE12 | $\Delta fimT$ | ADP1 *comP*<sup>T129C</sup> $\Delta fimT::gent$ | 1B, 1C, 1D, 2D, S1 | This study |
| CKE13 | $\Delta fimT + fimT$ | ADP1 *comP*<sup>T129C</sup> $\Delta fimT \Delta vanAB::kan$, *P*<sub>tac</sub>-*fimT* | 1B, 2D, 2E, 2F, S1 | This study |
| CKE14 | $\Delta fimT + fimT$-3xFLAG | ADP1 *comP*<sup>T129C</sup> $\Delta fimT::gent \Delta vanAB::kan$, *P*<sub>tac</sub>-*fimT*-3xFLAG | S1 | This study |
| CKE15 | $\Delta fimL \Delta pilT$ | ADP1 *comP*<sup>T129C</sup> $\Delta fimT::gent \Delta vanAB::kan$, *P*<sub>tac</sub>-*fimT*-3xFLAG $\Delta fimL::chlor \Delta pilT::spec$ | 2A, 2B | This study |
| CKE16 | Parent | ADP1 *comP*<sup>T129C</sup> $\Delta fimT::gent \Delta vanAB::kan$, *P*<sub>tac</sub>-*fimT* $\Delta pilT::spec$ | S3, 3A | This study |
| CKE17 | FimT-3xFLAG | ADP1 *comP*<sup>T129C</sup> $\Delta fimT::gent \Delta vanAB::kan$, *P*<sub>tac</sub>-*fimT*-3xFLAG $\Delta pilT::spec$ | S3 | This study |
| CKE18 | FimTR94Q-3xFLAG | ADP1 *comP*<sup>T129C</sup> $\Delta fimT::gent \Delta vanAB::kan$, *P*<sub>tac</sub>-*fimT*<sup>R94Q</sup>-3xFLAG $\Delta pilT::spec$ | S3 | This study |
| CKE19 | FimTK129Q-3xFLAG | ADP1 *comP*<sup>T129C</sup> $\Delta fimT::gent \Delta vanAB::kan$, *P*<sub>tac</sub>-*fimT*<sup>K129Q</sup>-3xFLAG $\Delta pilT::spec$ | S3 | This study |
| CKE20 | FimTR158QR163Q-3xFLAG | ADP1 *comP*<sup>T129C</sup> $\Delta fimT::gent \Delta vanAB::kan$, *P*<sub>tac</sub>-*fimT*<sup>R158QR163Q</sup>-3xFLAG $\Delta pilT::spec$ | S3 | This study |
| CKE21 | FimTR94QR158QR163Q-3xFLAG | ADP1 *comP*<sup>T129C</sup> $\Delta fimT::gent \Delta vanAB::kan$, *P*<sub>tac</sub>-*fimT*<sup>R94QR158QR163Q</sup>-3xFLAG $\Delta pilT::spec$ | S3 | This study |
| CKE22 | FimTK129QR158QR163Q-3xFLAG | ADP1 *comP*<sup>T129C</sup> $\Delta fimT::gent \Delta vanAB::kan$, *P*<sub>tac</sub>-*fimT*<sup>K129QR158QR163Q</sup>-3xFLAG $\Delta pilT::spec$ | S3 | This study |
| CKE23 | R33A | ADP1 *comP*<sup>T129C</sup> $\Delta fimT::gent \Delta vanAB::kan$, *P*<sub>tac</sub>-*fimT*<sup>R33A</sup> | 2D | This study |
| CKE24 | R41A | ADP1 *comP*<sup>T129C</sup> $\Delta fimT::gent \Delta vanAB::kan$, *P*<sub>tac</sub>-*fimT*<sup>R41A</sup> | 2D | This study |
| CKE25 | K53A | ADP1 *comP*<sup>T129C</sup> $\Delta fimT::gent \Delta vanAB::kan$, *P*<sub>tac</sub>-*fimT*<sup>K53A</sup> | 2D | This study |
| CKE26 | K55A | ADP1 *comP*<sup>T129C</sup> $\Delta fimT::gent \Delta vanAB::kan$, *P*<sub>tac</sub>-*fimT*<sup>K55A</sup> | 2D | This study |
| CKE27 | R62A | ADP1 *comP*<sup>T129C</sup> $\Delta fimT::gent \Delta vanAB::kan$, *P*<sub>tac</sub>-*fimT*<sup>R62A</sup> | 2D | This study |
| CKE28 | R90A | ADP1 *comP*<sup>T129C</sup> $\Delta fimT::gent \Delta vanAB::kan$, *P*<sub>tac</sub>-*fimT*<sup>R90A</sup> | 2D | This study |
| CKE29 | R92A | ADP1 *comP*<sup>T129C</sup> Δ*fimT::gent*<br>Δ*vanAB::kan*, *P*<sub>tac</sub>-*fimT*<sup>R92A</sup> | 2D | This study |
| CKE30 | R94A | ADP1 *comP*<sup>T129C</sup> Δ*fimT::gent*<br>Δ*vanAB::kan*, *P*<sub>tac</sub>-*fimT*<sup>R94A</sup> | 2D | This study |
| CKE31 | R96A | ADP1 *comP*<sup>T129C</sup> Δ*fimT::gent*<br>Δ*vanAB::kan*, *P*<sub>tac</sub>-*fimT*<sup>R96A</sup> | 2D | This study |
| CKE32 | K101A | ADP1 *comP*<sup>T129C</sup> Δ*fimT::gent*<br>Δ*vanAB::kan*, *P*<sub>tac</sub>-*fimT*<sup>K101A</sup> | 2D | This study |
| CKE33 | K111A | ADP1 *comP*<sup>T129C</sup> Δ*fimT::gent*<br>Δ*vanAB::kan*, *P*<sub>tac</sub>-*fimT*<sup>K111A</sup> | 2D | This study |
| CKE34 | R118A | ADP1 *comP*<sup>T129C</sup> Δ*fimT::gent*<br>Δ*vanAB::kan*, *P*<sub>tac</sub>-*fimT*<sup>R118A</sup> | 2D | This study |
| CKE35 | K129A | ADP1 *comP*<sup>T129C</sup> Δ*fimT::gent*<br>Δ*vanAB::kan*, *P*<sub>tac</sub>-*fimT*<sup>K129A</sup> | 2D | This study |
| CKE36 | K148A | ADP1 *comP*<sup>T129C</sup> Δ*fimT::gent*<br>Δ*vanAB::kan*, *P*<sub>tac</sub>-*fimT*<sup>K148A</sup> | 2D | This study |
| CKE37 | K152A | ADP1 *comP*<sup>T129C</sup> Δ*fimT::gent*<br>Δ*vanAB::kan*, *P*<sub>tac</sub>-*fimT*<sup>K152A</sup> | 2D | This study |
| CKE38 | K153A | ADP1 *comP*<sup>T129C</sup> Δ*fimT::gent*<br>Δ*vanAB::kan*, *P*<sub>tac</sub>-*fimT*<sup>K153A</sup> | 2D | This study |
| CKE39 | R158A | ADP1 *comP*<sup>T129C</sup> Δ*fimT::gent*<br>Δ*vanAB::kan*, *P*<sub>tac</sub>-*fimT*<sup>R158A</sup> | 2D | This study |
| CKE40 | R163A | ADP1 *comP*<sup>T129C</sup> Δ*fimT::gent*<br>Δ*vanAB::kan*, *P*<sub>tac</sub>-*fimT*<sup>R163A</sup> | 2D | This study |
| CKE41 | R94Q | ADP1 *comP*<sup>T129C</sup> Δ*fimT::gent*<br>Δ*vanAB::kan*, *P*<sub>tac</sub>-*fimT*<sup>R94Q</sup> | 2E, 2F | This study |
| CKE42 | K129Q | ADP1 *comP*<sup>T129C</sup> Δ*fimT::gent*<br>Δ*vanAB::kan*, *P*<sub>tac</sub>-*fimT*<sup>K129Q</sup> | 2E, 2F | This study |
| CKE43 | R158QR163Q | ADP1 *comP*<sup>T129C</sup> Δ*fimT::gent*<br>Δ*vanAB::kan*, *P*<sub>tac</sub>-*fimT*<sup>R158QR163Q</sup> | 2E, 2F | This study |
| CKE44 | R94QR158QR163Q | ADP1 *comP*<sup>T129C</sup> Δ*fimT::gent*<br>Δ*vanAB::kan*, *P*<sub>tac</sub>-*fimT*<sup>R94QR158QR163Q</sup> | 2E, 2F | This study |
| CKE45 | K129QR158QR163Q | ADP1 *comP*<sup>T129C</sup> Δ*fimT::gent*<br>Δ*vanAB::kan*, *P*<sub>tac</sub>-*fimT*<sup>K129QR158QR163Q</sup> | 2E, 2F | This study |
| CKE46 | - | ADP1 *comP*<sup>T129C</sup> Δ*fimT::gent*<br>Δ*pilT::spec* | 3A | This study |
| CKE47 | R94QR158QR163Q | ADP1 *comP*<sup>T129C</sup> Δ*fimT::gent*<br>Δ*vanAB::kan*, *P*<sub>tac</sub>-*fimT*<sup>R94QR158QR163Q</sup><br>Δ*pilT::spec* | 3A | This study |
| CKE48 | R94Q | ADP1 *comP*<sup>T129C</sup> Δ*fimT::gent*<br>Δ*vanAB::kan*, *P*<sub>tac</sub>-*fimT*<sup>R94Q</sup> Δ*pilT::spec* | 3A | This study |
| CKE49 | K129Q | ADP1 *comP*<sup>T129C</sup> Δ*fimT::gent*<br>Δ*vanAB::kan*, *P*<sub>tac</sub>-*fimT*<sup>K129Q</sup><br>Δ*pilT::spec* | 3A | This study |
| CKE50 | R158QR163Q | ADP1 *comP*<sup>T129C</sup> Δ*fimT::gent*<br>Δ*vanAB::kan*, *P*<sub>tac</sub>-*fimT*<sup>R158QR163Q</sup><br>Δ*pilT::spec* | 3A | This study |
| CKE51 | K129QR158QR163Q | ADP1 *comP*<sup>T129C</sup> Δ*fimT::gent*<br>Δ*vanAB::kan*, *P*<sub>tac</sub>-*fimT*<sup>K129QR158QR163Q</sup><br>Δ*pilT::spec* | 3A | This study |
| CE317 | | ADP1 *comP*<sup>T129C</sup> Δ*pilT::spec* | 3C | <sup>1</sup> |
| CKE52 | Parent or WT | *Acinetobacter nosocomialis* M2 | 4D, S5 | <sup>2</sup> |
| CKE53 | *pilA::apr*, *pilA* | M2 *pilA::apr*, *pilA* | S5 | This study |
| CKE54 | *pilA::apr*, *pilA*<sup>S77C</sup> | M2 *pilA::apr*, *pilA*<sup>S77C</sup> | S5 | This study |
| CKE55 | Parent | M2 *pilA::kan*, *pilA*<sup>S77C</sup> | 4B, 4C | This study |
| CKE56 | Δ*fimT* | M2 *pilA::kan*, *pilA*<sup>S77C</sup> Δ*fimT::spec* | 4B, 4C | This study |
| CKE57 | Δ*fimT* | M2 Δ*fimT::spec* | 4D | This study |
| CKE58 | Δ*fimT* *P*<sub>tac</sub>-*fimT*<sup>An</sup> | M2 Δ*fimT::spec* Δ*vanAB::kan*, *P*<sub>tac</sub>-*fimT*<sup>An</sup> | 4D | This study |
| CKE59 | Δ*fimT* *P*<sub>tac</sub>-*fimT*<sup>Ab</sup> | M2 Δ*fimT::spec* Δ*vanAB::kan*, *P*<sub>tac</sub>-*fimT*<sup>Ab</sup> | 4D | This study |

**Table S2.**
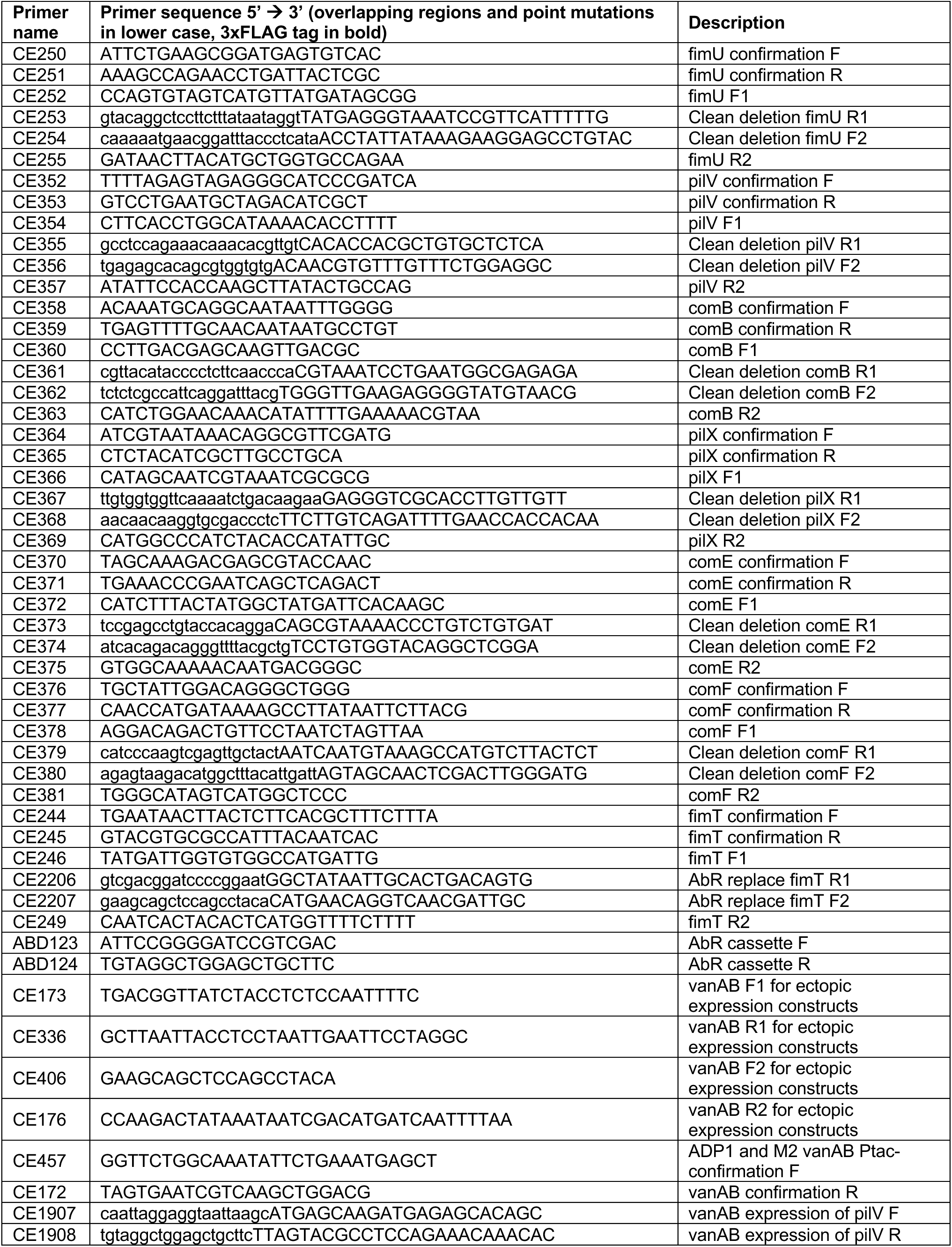
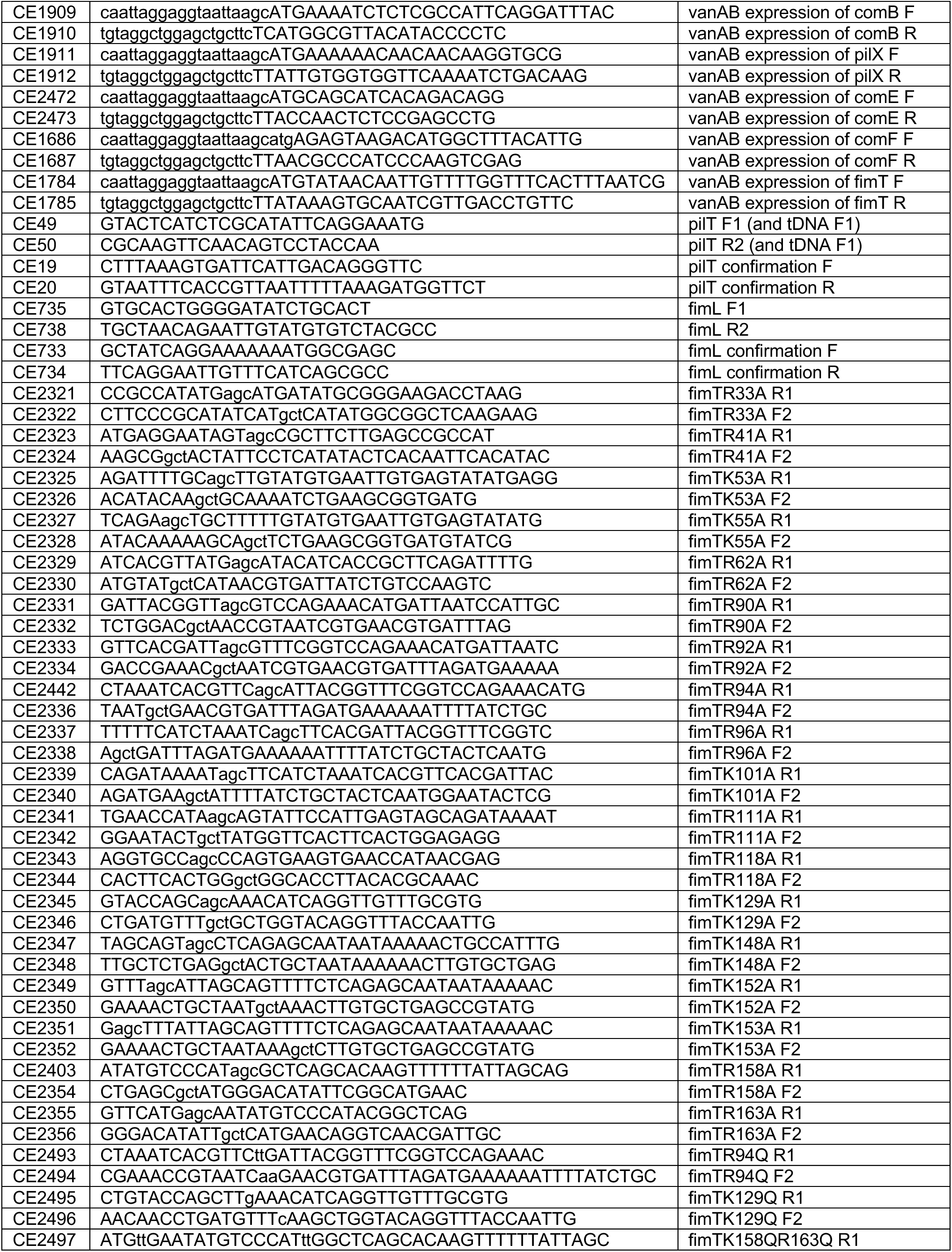
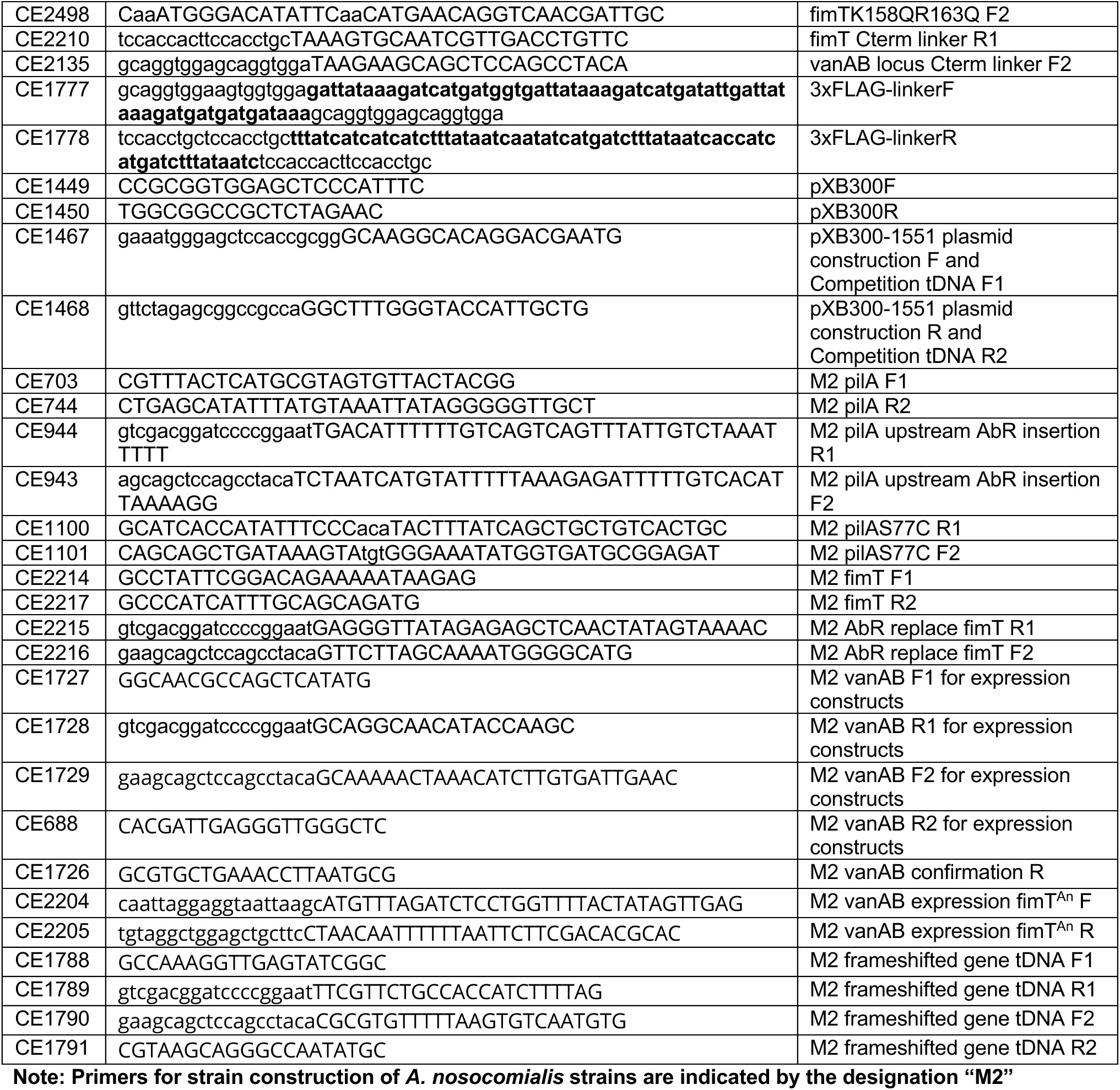
Primers used for strain construction and confirmation.

